# A Subset of HIV-1 Controllers Lack Cortical Actin Disruption Indicative of ARP2/3 Inhibition

**DOI:** 10.1101/2023.07.27.550860

**Authors:** Robert L. Furler O’Brien, Colin Kovacs

## Abstract

A small fraction of people living with HIV-1 suppress viral replication naturally and exhibit delayed or absent disease progression without antiretroviral therapy, yet the underlying mechanisms of viral control remain elusive. Despite the known role of HIV-1 in disrupting the actin cytoskeleton and altering cell migration and morphology within tissues, the molecular underpinnings that link viral actin disruption to disease progression have yet been linked to disease progression. We have previously shown through ultrastructural and time-lapse imaging that HIV-1 mediated actin disruption mirrors ARP2/3 inhibition within primary CD4^+^ T cells of normal progressors and uninfected controls. Infected CD4^+^ T cells from these two cohorts routinely exhibit two unique phenotypes when migrating. The first morphological difference is a sharp elongated and pointed lamellipodial tip, “Rhino” phenotype, distinct from the broad leading edge of uninfected cells. The second morphological difference is a non-apoptotic polarized blebbing at the lamellipodia of infected cells. These two pathological morphologies can be recapitulated in uninfected cells with chemical inhibitors of the ARP2/3 complex and are temporally linked based on the differentiation status of the T cell. These effects are dampened, but not totally eradicated, in the absence of the HIV-1 Nef protein. In contrast to normal progressors, infected cells from two out of the three HIV-1 controllers tested in this study did not exhibit these cellular pathologies. The profound impact of ARP2/3 inhibition on immunopathogenesis within genetic and infectious diseases provides context into how HIV-1 may cause cellular and systemic immune dysfunction in normal progressors. The mechanically destabilized cellular cortex may also provide a selective protection for viral genome-intact and long-lived defective reservoirs from cell-mediated killing by host CD8^+^ T cells and NK cells. This mechanical instability is absent in some HIV-1 controllers. Restoring ARP2/3 function and cortical actin integrity in people living with HIV-1 infection is a new avenue of investigation to eradicate HIV-1 infected cells from the body.

---

After four decades of intense study, how some individuals control HIV-1 disease progression without antiretroviral therapy (ART) is still largely unknown. Since the beginning of the HIV-1 pandemic, reports of virological and immunological HIV-1 controllers have suggested there are multiple mechanisms of disease control including both viral and host determinants^1,2^. However, in the early 1990s, cohorts of patients with delayed disease progression began to solidify the link between the viral Nef protein and Acquired Immunodeficiency Syndrome (AIDS). In 1992, Learmont *et al*. described a cohort in New South Wales consisting of one HIV-1 infected individual and 6 recipients of that individual’s blood or blood products who subsequently became infected but showed few signs of AIDS progression or CD4^+^ T cell decline even after 10.75-14 years following infection in the absence of ART^3-5^. Viral sequence analysis revealed a common deletion in the *nef* gene and the overlapping U3 region of the LTR, while other HIV-1 genes remained intact^3^. This cohort became known as the Sydney Blood Bank Cohort and was one of the first reports that AIDS progression could be stunted or completely inhibited in humans due to mutations in the *nef* gene.

Although deletions or functional mutations in the *nef* gene may lead to HIV-1 control, other undefined mechanisms exist. A study of 12 HIV-1 infected Kenyan children with slow disease progression showed no detectable deletion or functional alteration in *nef* or other viral genes, suggesting underlying host-mediated mechanisms of control exist in absence of viral mutations^6^. The majority of HIV-1 controllers are infected with fully-intact replication competent virus and control may be linked to host genetic mutations that hinder HIV-1 entry, limit high viral replication, or promote effective viral clearance.

Genome wide association studies (GWAS) were undertaken to pinpoint genetic contributions to HIV-1 control and have been reviewed elsewhere^7,8^. The major genetic findings of over a dozen GWAS since 2007 have linked disease progression and acquisition to mutations in chemokine receptors and specific alleles within the HLA loci of the human genome including HLA-B57. Although certain HLA alleles may be increased in HIV-1 controller cohorts, these specific alleles are also found in normal progressor cohorts, suggesting HLA alleles alone do not guarantee control. Despite multiple GWAS studies, no group has been able to make reproducibly significant predictions of genetic mutations that can mechanistically explain HIV-1 control at the biological level other than HIV-1 coreceptor mutation (CCR5Δ32). This only known host mutation that protects from HIV-1 entry and disease progression is an internal deletion in the *CCR5* gene, whose protein product is a co-receptor for HIV-1 entry. This *CCR5Δ32* mutation is rare, and homozygosity is required for full protection against R5-tropic isolates of HIV-1. One potential caveat of these GWAS studies is the analyses of single nucleotide polymorphisms (SNPs) that may not be associated with larger insertions, deletions, or mutations that may exist within the controller populations. Additionally, if multiple mechanisms of HIV-1 control exist, genetic components may be missed with these standard methodologies.

Aside from genetic studies, several *in vivo* and clinical studies show that a robust antiviral CD8^+^ T cell effector function is key to delaying or preventing disease progression in many controllers^9,10^. Integration site analysis of HIV-1 infected cells from controllers shows an increased propensity of provirus integration within heterochromatic regions, suggesting a selective removal of cells where virus was integrated into euchromatic regions and could be expressed^11^. This may be indicative of a robust CD8^+^ T cell response to cells with active viral replication. However, it is uncertain if HIV-1 controllers are capable of enhanced anti-viral CD8^+^ T cell responses or if their infected CD4^+^ T cells are inherently susceptible to targeting and elimination.

In this study, the latter hypothesis is supported with new findings that HIV-1 infected CD4^+^ T cells from controllers exhibit mechanical characteristics of target cells that allow for effective CTL clearance^12,13^. In contrast, infected cells from normal progressors and healthy donors have disrupted cortical actin structures and resemble leukocytes in patients with primary immunodeficiencies that stem from mutations in actin regulatory genes, specifically those that affect the ARP2/3 complex. Furthermore, these HIV-1 induced actinopathies are partially but not completely due to HIV-1 Nef, a known viral determinant of disease progression. A link between cortical actin integrity in HIV-1 infected cells and their susceptibility to CTL-mediated elimination remains to be tested; however, this striking contrast in HIV-1 infected CD4^+^ T cell morphology between normal progressors and some HIV-1 controllers provides new evidence that an underlying mechanical susceptibility of infected cell elimination may exist in HIV-1 controllers. Restoring cortical actin integrity in normal progressors may be key to the effective elimination of HIV-1 infected cells from the body.

Evidence that these newly identified HIV-1 induced morphological alterations found in normal progressors and healthy controls influence disease progression is still lacking. In this study, we characterized the ability of HIV-1 to alter cortical actin in three HIV-1 controllers, each with a history of ART-free viral suppression (**Table 1 and Extended Data Table 1)**.

**Table 1.** Summary of HIV Controller Demographics and Clinical Parameters.

| Item | HIV Controller 1 | HIV Controller 2 | HIV Controller 3 |
| --- | --- | --- | --- |
| Demographics |  |  |  |
| Age (Years) | 78 | 53 | 56 |
| Sex | M | M | M |
| Race | Caucasian | Caucasian | Caucasian |
| Clinical Information |  |  |  |
| Duration of Infection (years) | 34 | 27 | 26 |
| Peak Viremia off ART (copies/mL) | 372 | 3182 | 55 |
| Absolute CD4+ T Cell Nadir (cells/mL) | 470 | 418 | 440 |
| Genetic Information |  |  |  |
| HLA B5701 Present | No | Yes | Yes |
| CCR5 Genotype (*wildtype* or *CCR5*Δ32) | *wt/wt* | *wt/wt* | *wt/Δ32* |
| Other |  |  |  |
| Participant in Previous Controller Study | Subject 4 | NA | Subject 3 |

CD4^+^ T cells from these controllers were readily infectable *in vitro*, similar to controllers from other reports. Using single-round pseudo-typed reporter viruses, Rabi *et al*. reported that unstimulated primary CD4^+^ T cells from controllers could be infected with HIV-1 as readily as cells from viremic or uninfected controls^14^. The impact of cell activation on infection susceptibility became more appreciated and others reported that although CD4^+^ T cells from Elite Controllers (EC) were readily infected, they produced less virions compared to cells from normal progressors^15^. To assess infectibility and characterize the morphological changes that occur in infected cells of HIV-1 controller cells, previously activated and expanded CD4^+^ T cells were infected with full length EGFP reporter viruses and allowed to migrate on fibronectin-coated dishes. As shown in **Figure 1 and Supplementary Video 1-2**, infected CD4^+^ T cells from two of the three HIV-1 controllers (**Figure 1A and 1B**) exhibit normal morphologies with broad lamellipodia and no polarized blebbing. Interestingly one of the controllers still retained rare cells with the Rhino phenotype despite an absence of polarized blebbing, suggesting there are likely donor differences in actin stability and control. The Rhino phenotype was not observed in any of the cells infected with ΔNef EGFP.

**Figure 1:**
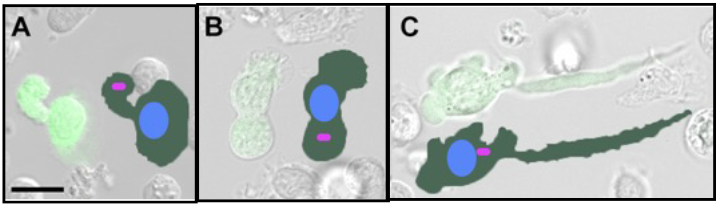
A subset of HIV-1 Controllers Retain Typical Morphologies Despite Infection. (A) HIV-1 Controller 1 and (B) Controller 2 retain broad lamellipodia and extended uropods in infected cells; however, (C) Controller 3 exhibited abnormal morphologies similar to cells from normal HIV-1 progressors and uninfected donors. Outlines of morphologies with respective location of nuclei (blue) and uropods (fuscia rods) shown for anatomical orientation. Scale = 10μm.

The third controller (**Figure 1C and Supplementary Video 3**) had a unique clinical history compared to the first two controllers (**Figure 2**) with undetectable viral loads for the majority of the time period and an absence of anti-HIV-1 specific CD8^+^ T cell responses that were previously measured^16^.

**Figure 2:**
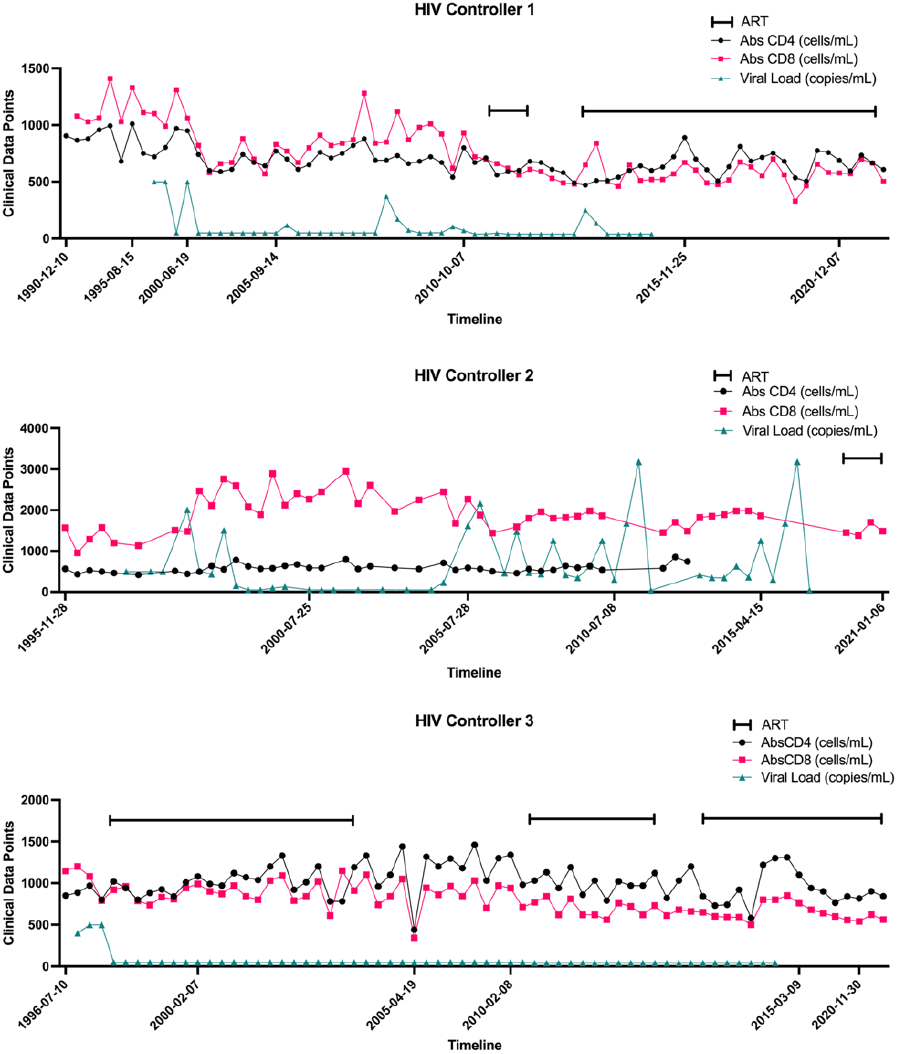
Clinical timeline of three HIV 1 controllers examined in this study. CD4+ T cells counts (black), CD8+ T cell counts (fuchsia), and viral loads (teal) are displayed over the study period. Periods of antiretroviral therapy are indicated by brackets. Expanded measurements can be found in Extended Data Table 1.

Following *in vitro* infection with a reporter virus, the infected cells from the third controller clearly exhibited the abnormal phenotypes seen in normal progressors. Genetic screening of the three HIV-1 controllers indicated that unlike the first two controllers, the third was heterozygous for the CCR5Δ32 mutation (**Extended Data Figure 1**) and has the HLA-B5701 allele. A recent report of an individual with a similar genotype (CCR5Δ32^+/-^ and HLA-B5701^+^), suggested possible elimination of infection with a *nef-*deleted virus^17^. Reservoir measurements and anti-HIV-1 specific CD8^+^ T cell responses were measured for Controllers 1 and 3 in a previous study^16^. Although *in vitro* quantitative viral outgrowth assays (QVOA) were done on Controller 3, viral sequencing was not done to confirm the presence of *nef* within any potential intact viruses found in this participant. We and others have shown that *nef-*deleted viruses can replicate easily during *in vitro* assays, including QVOA. The additional lack of anti-HIV-1 specific CD8^+^ T cell responses in Controller 3 and the phenotypic differences we observed in this study may indicate a possible separate mechanism of control in Controller 3 versus Controllers 1 and 2. The clear distinction between the phenotypes of Controllers 1 and 2 to Controller 3 and normal progressors suggest microscopic evaluation may be useful in classifying different types of HIV-1 control.

This and previous studies indicate that actin destabilization is not required for ongoing HIV-1 replication *in vitro* or *in vivo*; however, the ability to retain stable actin structures in infected cells may be a key factor in controlling disease progression. HIV-1 replicates within CD4^+^ T cells from HIV-1 controllers *in vitro* and *in vivo*, although they may produce less virions than normal progressors^15^. Although Nef has clearly been linked to disease progression, Nef has many functions but its actin disrupting ability has not been directly linked to disease progression. We provide evidence that some HIV-1 controllers may thwart Nef’s ability to disrupt cortical actin. This along with recent *in vivo* studies begins to provide a new model of HIV-1 control that needs further scrutiny. In a viral kinetics study using a humanized mouse model, HIV-1 Nef was reported to disrupt the actin cytoskeleton within infected CD4^+^ T cells and reduce cell migration speed by half^18^. This effect was mapped to Nef residue F191A, a critical residue for PAK2 signaling. Interestingly, viral competition studies in humanized mice showed that Nef-deleted (ΔNef) viruses propagated much faster in the first few weeks of infection indicating a selective disadvantage for intact Nef *in vivo*. However, in later timepoints the wild-type strain containing Nef predominated, indicating a switch in selection from ΔNef to wild-type Nef viruses around the time when CD8^+^ T cell numbers were increasing. This potential immune evasion may initially be correlated to Nef’s ability to downregulate MHC-I; however, the F195A mutant used in this study has not been reported to affect this function of Nef. The cortical actin destabilization caused by wild-type Nef may hinder effective CD8^+^ T cell responses. This hypothesis needs further evaluation.

Intact ARP2/3 and cortical actin within HIV-1 contollers may allow for more efficient CD8^+^ T cell targeting. Recently, a correlation between cytoskeletal rigidity in cancer cells and their ability to be targeted by cytotoxic T lymphocytes in mice highlighted the need for stable actin structures in target cells during CTL-mediated killing^12,13^. Destabilized cortical actin structures within infected cells may protect the tumor or infected cells from CTL-mediated attack in 3-dimensional environments or delay the kinetics of attack.

If one mechanism of HIV-1 control is the ability to retain ARP2/3 function and cortical actin integrity following infection, many HIV-1 cure strategies may not overcome this barrier alone. Passive immunization of broadly neutralizing antibodies and immune-stimulating biologics or infusion of HIV-1 specific effector cells will unlikely eradicate or robustly suppress viral replication long term without addressing the inherent HIV-1 induced cortical actin disruption found in normal progressors. However, restoring cortical actin integrity within infected cells may lead to more effective clearance of both the fully-intact and the defective persistent HIV-1 reservoirs by the host’s natural immune system.

## Online Methods

### Study Participants

Small volume whole blood draws or frozen PBMC samples from deidentified HIV-1 controllers, normal progressors, and healthy participants were obtained from the Maple Leaf Clinic (Toronto, Canada) and Gulf Coast Blood Center (Houston, Texas, USA). All participants signed informed-consent forms approved by their respective Investigational Review Boards (IRB).

### CD4^+^ T cell isolation and activation

Freshly acquired blood was processed by Ficoll-Paque separation to isolate peripheral blood mononuclear cells (PBMC) from healthy donors, normal HIV-1 progressors, and HIV-1 controllers. Negative selection of total CD4^+^ T cells was done using StemCell EasySep Human CD4^+^ T Cell Enrichment Kit (Catalog #19052). Cells were than cultured in RPMI, 10% fetal bovine serum (FBS), penicillin, streptomycin, and glutamine at one to two million cells/mL. Infections were done on negatively selected total human CD4^+^ T cells that were previously activated with anti-CD3 (Ultra-LEAF™ Purified anti-human CD3 Antibody, clone OKT3, Biolegend, Catalog #317325) and anti-CD28 (Ultra-LEAF™ Purified anti-human CD28 Antibody, Clone CD28.2, Biolegend, Catalog #302933) antibodies in the presence of rhIL-2 and IL-15. The media was changed every three to four days for two weeks prior to infection. Post-activation and prior to infection, a second round of negative selection was done to achieve 97-99% CD4^+^ T cell purity as measured by flow cytometry. The cells were then resuspended at a concentration of 2*10^7^ cells/mL in RPMI supplemented with 10% FBS, penicillin, streptomycin, glutamine, IL-2 and IL-15 prior to infection.

### Infection

For each of the infection and mock conditions, 500uL of cells were added to a well in a 6-well plate before adding 500uL of virus or mock supernatant. The cell/virus mixture was spinoculated at 2900 rpm (1965g) for 2 hours at 37°C. These cells were added to an equal number of non-spinoculated cells and cultured at 2*10^6^ cells/mL at 37°C for up to 3 weeks, with microscopic analysis and media changes occurring every 2-3 days. To optionally increase the population of infected cells, cells were crowded at Day 3 post-infection by culturing in 200uL at 2*10^6^ cells/mL in 96-well ‘U’-bottom plates for an additional 3 to 4 days. Monitoring of infected cells populations was done by intracellular HIV-1 Gag p24 measurements using flow cytometry and measuring EGFP fluorescence by microscopy when using reporter viruses.

Cells were infected with primary, laboratory, and reporter isolates of CCR5-tropic or CCR5/CXCR4 dual-tropic Human Immunodeficiency Virus-1. The following three viruses were obtained from the NIH AIDS Reagents program: HIV-1 ADA (ARP-416, CCR5-tropic), HIV-1 92/BR/014 (ARP-1753, Dual-tropic, syncytium inducing), HIV-1 92/BR/004, ARP-1752 (CCR5-tropic, non-syncytium inducing). Reporter viruses were generated from the following plasmids that were generously provided by Dr. Thorsten Mempel and Dr. Shariq Usmani: HIV-1 (WT) IRES-EGFP (pBR43IeG nef+ R-BaL env) and HIV-1 (ΔNef) IRES-EGFP (pBR43IeG nef+ R-BaL env, Nef deleted)^18^. Plasmids were transformed into Stbl2 competent *E. coli* before clonal selection and viral sequencing confirmation. Confirmed clones were transfected into HEK 293 cells to produce virions that were then filtered and quantified by p24 ELISA and TZM-bl assay.

### CCR5Δ32 Screening

Genetic screening for the known protective variant (CCR5Δ32) of the HIV-1 co-receptor CCR5 was done as previously described^19^. Briefly, genomic DNA was extracted from 5*10^6^ PBMC, using Qiagen DNeasy Blood & Tissue Kit (Catalog #69506). A segment of the *CCR5* gene was amplified from each study participant by PCR in standard conditions using the following primer pair: CCR5-F: 5’-CTCCCAGGAATCATCTTTACC-3’ and CCR5-R: 5’-TCATTTCGACACCGAAGCAG-3’. PCR amplicons were separated on a 2% agarose gel at 130V for 60 minutes before imaging on a Bio-Rad Chemidoc workstation. A PCR fragment of 200 bp indicates a wild-type CCR5 allele while a fragment of 168 bp indicates a CCR5Δ32 allele.

### Time Lapse Confocal Microscopy

Time lapse confocal imaging was done in an incubated chamber. Image frames were taken at 1 second intervals with either 25X, 40X, or 63X objectives. Videos were generated using a speed rate of 25 frames per second using Zeiss ZenBlue Software. When tracking organelle location using fluorescent dyes, 1μM MitoTracker™ Deep Red FM (ThermoFisher, Catalog number: M22426) was used to detect mitochondria and 1μM LysoTracker™ Blue DND-22 (ThermoFisher, Catalog number: L7525) was used to detect acidic organelles including lysosomes and nuclei.

## Acknowledgements

The following reagents were obtained through the NIH HIV Reagent Program, Division of AIDS, NIAID, NIH: HIV-1 ADA (ARP-416), contributed by Dr. Howard Gendelman; HIV-1 92/BR/014 (ARP-1753) and HIV-1 92/BR/004 (ARP-1752), contributed by UNAIDS Network for HIV Isolation and Characterization; and Human rhIL-2 from Dr. Maurice Gately, Hoffmann -La Roche Inc. (ARP-136). Human IL-15 was obtained through the Frederick National Laboratory for Cancer Research Biological Resources Branch pre-clinical repository. The following plasmids were generously provided by Dr. Thorsten Mempel and Dr. Shariq Usmani: HIV=1 (WT) IRES-EGFP (pBR43IeG nef+ R-BaL env), HIV-1 (ΔNef) IRES-EGFP (pBR43IeG nef+ R-BaL env, Nef deleted)^18^. Sample procurement was completed with the help of Erika Benko. This publication resulted in part from research supported by National Institute of Allergy and Infectious Diseases (NIAID) awards UM1 AI126617, co-funded by National Institute on Drug Abuse (NIDA), National Institute of Mental Health (NIMH), and National Institute of Neurological Disorders and Stroke (NINDS).

## Author contributions

CK performed the clinical evaluation and procurement of the HIV-1 positive samples. RLFO performed the electron and light microscopy. The conception of the work, the interpretation of results, and manuscript preparation was done by RLFO. All authors reviewed and edited the final manuscript.

## Materials & Correspondence

Supplementary Information is available for this paper. Correspondence and requests for materials should be addressed to Robert Furler O’Brien at. Reprints and permissions information is available at www.nature.com/reprints.

## Supplementary information

### Extended Data

**Extended Data Figure 1:**
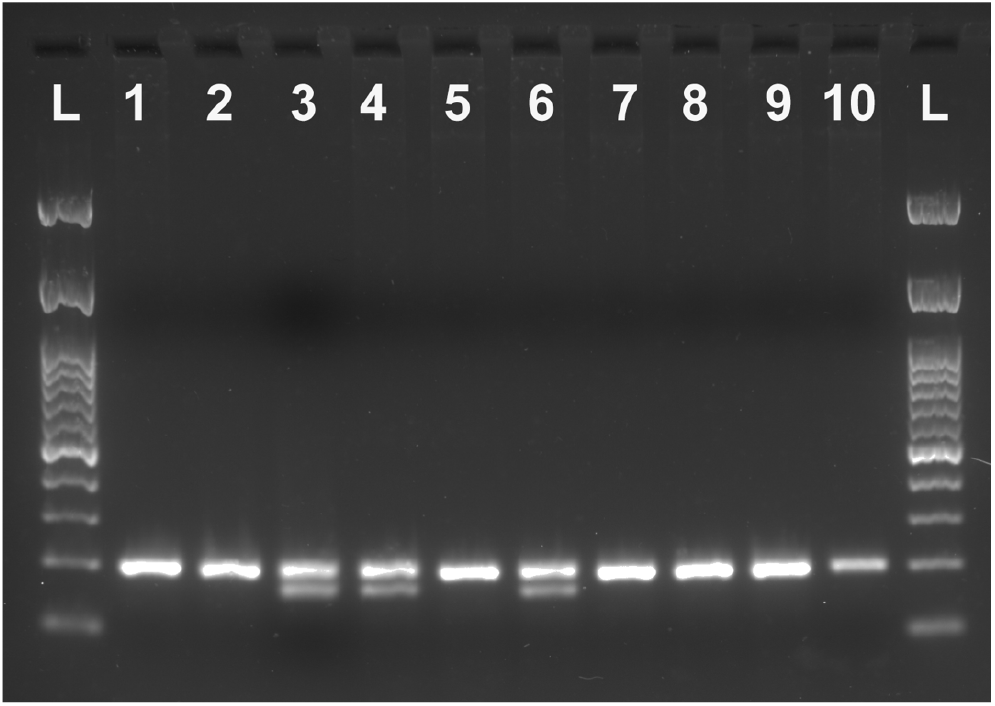
CCR5 Genotyping of HIV-1 Controllers and Progressors in this study. A segment of the *CCR5* gene was amplified to detect the presence or absence of the CCR5Δ32 allele in both HIV-1 Controllers and Normal Progressors. A higher band in each lane shows a PCR fragment of 200 bp, which indicates a wild-type CCR5 allele. A lower band present in some lanes show a PCR fragment of 168 bp, which indicates a CCR5Δ32 allele. HIV-1 Controller 1 and 2 (Lanes 1 and 2 respectively) do not carry a CCR5Δ32 allele while HIV-1 Controller 3 is heterozygous for the CCR5Δ32 allele.

Extended Data Table 1 – Excel spreadsheet of clinical parameters for HIV-1 positive normal progressors and controllers in this study.

Supplementary Video 1: HIV-1 Controller 1 cells infected with HIV-1 WT EGFP^+^, (time-lapse).

Supplementary Video 2: HIV-1 Controller 2 cells infected with HIV-1 WT EGFP^+^, (time-lapse).

Supplementary Video 3: HIV-1 Controller 3 cells infected with HIV-1 WT EGFP^+^, (time-lapse).

## Data availability

The data presented in this study are available in the supplementary material.

## Life sciences and behavioural & social sciences reporting guidelines

### Human subject data

Deidentified biospecimens were used in these studies. All samples were obtained under institutional IRB approval with documented informed consent.

